# PARP10 is Critical for Stress Granule Initiation

**DOI:** 10.1101/2023.10.13.562236

**Authors:** Aravinth Kumar Jayabalan, Krishna Bhambhani, Anthony K L Leung

**Affiliations:** Department of Biochemistry and Molecular Biology, Bloomberg School of Public Health, Johns Hopkins University, Baltimore, MD 21205, USA; Department of Chemical and Biomolecular Engineering, Whiting School of Engineering, Johns Hopkins University, Baltimore, MD 21205, USA; McKusick-Nathans Department of Genetics Medicine, School of Medicine, Johns Hopkins University, Baltimore, MD 21205, USA; Department of Oncology, School of Medicine, Johns Hopkins University, Baltimore, MD 21205, USA; Department of Molecular Biology and Genetics, School of Medicine, Johns Hopkins University, Baltimore, MD 21205, USA

## Abstract

Stress granules (SGs) are cytoplasmic biomolecular condensates enriched with RNA, translation factors, and other proteins. They form in response to stress and are implicated in various diseased states including viral infection, tumorigenesis, and neurodegeneration. Understanding the mechanism of SG assembly, particularly its initiation, offers potential therapeutic avenues. Although ADP-ribosylation plays a key role in SG assembly, and one of its key forms—poly(ADP-ribose) or PAR—is critical for recruiting proteins to SGs, the specific enzyme responsible remains unidentified. Here, we systematically knock down the human ADP-ribosyltransferase family and identify PARP10 as pivotal for SG assembly. Live-cell imaging reveals PARP10’s crucial role in regulating initial assembly kinetics. Further, we pinpoint the core SG component, G3BP1, as a PARP10 substrate and find that PARP10 regulates SG assembly driven by both G3BP1 and its modeled mechanism. Intriguingly, while PARP10 only adds a single ADP-ribose unit to proteins, G3BP1 is PARylated, suggesting its potential role as a scaffold for protein recruitment. PARP10 knockdown alters the SG core composition, notably decreasing translation factor presence. Based on our findings, we propose a model in which ADP-ribosylation acts as a rate-limiting step, initiating the formation of this RNA-enriched condensate.

**HIGHLIGHTS:**

- PARP10 plays a crucial role in the initial SG assembly kinetics.
- The core SG component G3BP1 is a substrate of PARP10.
- PARP10 is required for SG assembly mediated by G3BP1 or its synthetic mimic.
- PARP10 knockdown reduces the levels of translation factors within the SG core.

## INTRODUCTION

Stress granules (SGs) are cytoplasmic condensates formed in response to various stressors, including viral infection and oxidative stress^1,2^. The assembly and disassembly of SGs are tightly regulated processes in response to and relief from stress, respectively^3^. Implicated in tumorigenesis, neurodegeneration, and antiviral innate immunity, elucidating the mechanisms underlying SG formation could offer new therapeutic avenues for various diseases^4–8^.

Upon stress, cells halt global translation, releasing mRNAs from ribosomes. These naked mRNAs interact and condense with themselves as well as with translation factors and RNA-binding proteins in the cytoplasm to form SGs^1^. Post-translational modifications further fine-tune this process, providing precise spatiotemporal control over SG assembly and disassembly^9–14^. Yet, the exact mechanisms that initiate SG assembly remain to be defined.

ADP-ribosylation—the addition of one or more ADP-ribose units onto target proteins—plays roles in diverse biological processes^15^, such as DNA damage repair, mRNA translation, protein degradation, and the formation of biomolecular condensates, including SGs^14,16^. Of the 17 ADP-ribosyltransferases in humans, commonly known as PARPs, all except PARP13 are catalytically active. Specifically, PARPs 1, 2, 5a, and 5b add multiple ADP-ribose units, forming poly(ADP-ribose) or PAR, while the remaining 11 add single units^15,17^, resulting in mono-ADP-ribosylation or MARylation^17^.

Within SGs, five PARPs (PARPs 5a, 12, 13, 14, 15) and PAR glycohydrolase (PARG) are identified. Overexpressing PARG suppresses SG formation, whereas the knockdown of this PAR-degrading enzyme impedes their disassembly^14^. Therefore, PAR is key for SG assembly, acting as a structural scaffold and facilitating protein recruitment to SGs^16,18–20^. Despite the importance of PAR in SG assembly, the role of MARylation remains elusive, even though three out of five PARPs localized in SGs add MAR only.

Intriguingly, overexpression of any of these PARP can induce SGs^14^. Our recent work further revealed that a conserved MAR-degrading enzyme in the non-structural protein 3 (nsP3) of alphaviruses is critical for viral replication, pathogenesis, and SG dynamics. While infection initially triggers SG formation, subsequent disassembly relies on nsP3’s MAR-degrading activity^18^.

Upon stress, RNA-binding proteins, including G3BP1, undergo increased ADP-ribosylation^14,21,22^. G3BP1 and its paralogs G3BP2 are central to SG assembly, and their absence renders cells unable to form SGs under various stresses^23^. Apart from being ADP-ribosylated, G3BPs can bind to PAR. Notably, during infection, the viral-encoded MAR-degrading nsP3 also removes ADP-ribosylation from G3BP1^18^. These findings underscore the importance of ADP-ribosylation of G3BP1 in SG formation.

Despite recognizing ADP-ribosylation’s crucial role in SG assembly, we still lack clarity on which PARP(s) regulate this process, their exact roles, and targets within SGs^14,16,18,21^. Addressing these gaps could pave the way for developing inhibitors to modulate SG formation therapeutically^6^. Through systematic knockdown, we identified PARP10—which adds MAR only—as critical for SG assembly. Our live-cell imaging demonstrated that diminished PARP10 levels delay SG formation, and our fractionation experiment showed reduced translation factor levels in the SG core upon PARP10 knockdown. We also pinpointed G3BP1 as a substrate targeted by PARP10. We conclude with a model proposing how MARylation regulates SG assembly kinetics, particularly during the initial stage.

## RESULTS

### MARylation is critical for SG assembly

Previously, we observed a notable difference in suppressing SG assembly by expressing hydrolase against PAR vs. MAR^18^. Human PARG degrades all but the last ADP-ribose conjugated to the protein^24^, thereby effectively converting PAR to MAR, while chikungunya nsP3 removes MAR from protein^25^. Compared with the PAR-degrading PARG, the MAR-degrading nsP3 has a stronger effect suppressing SG assembly^18^. These findings led us to hypothesize that MARylation is critical for SG assembly.

To investigate further, we co-expressed MAR and PAR hydrolases at different expression levels and assessed their impact on SG assembly. First, we maintained the expression of the PAR-degrading enzyme PARG constant while gradually increasing the expression of MAR-degrading nsP3. Typically, arsenite treatment for 30 min induced SG assembly in ∼90% of U2OS cells; PARG alone inhibited its assembly, resulting in ∼60% of cells with SGs, as indicated by the diffused distribution of the marker eIF3b (Fig. 1A, 1B, red bars). However, the co-expression of nsP3 significantly extended the inhibition range, resulting in ∼70% of cells without SGs (Fig. 1A-C). Notably, even adding the least amount of nsP3 had an obvious effect on SG inhibition. Conversely, when nsP3 was expressed alone, ∼70% of cells were without SGs, whereas further addition of PARG only had a mild effect on SG inhibition (Fig. 1A, 1B, green bars). These findings collectively suggest that the removal of MARylation is crucial for inhibiting SG assembly.

**Figure 1.**
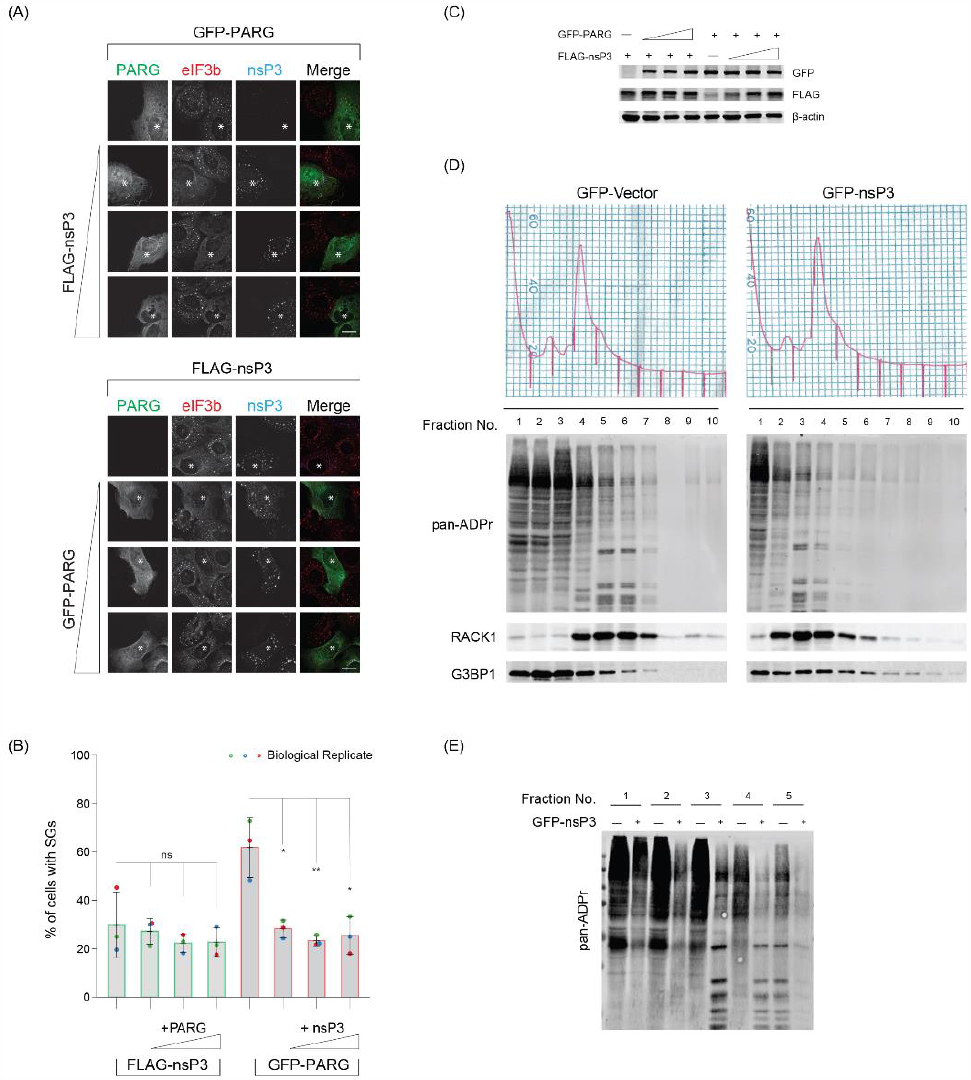
Removal of terminal ADP-ribose is critical to inhibit SGs. (A) U2OS cells were transfected with FLAG-tagged nsP3 alone or co-transfected with increasing concentrations of GFP-tagged PARG (Bottom panel) or GFP-tagged PARG alone or with increasing concentrations of FLAG-tagged nsP3 (Top panel). 24 h post transfection, cells were treated with 0.2 mM arsenite for 30 min, fixed, and immunostained against eIF3b and FLAG antibodies. Scale bar 10 μm. Asterisk indicates co-transfected cells. Scale bar, 10 μm. (B) Bar graph showing the percentage of GFP/FLAG positive cells containing SGs in panel A. Cells were scored as SG positive when eIF3b condensates are present in cells expressing both FLAG- and GFP-tagged constructs. (C) Western blot data confirming the expression of transfected constructs in panel A. (D) U2OS cells either transfected with GFP or GFP-nsP3 were treated with 0.2 mM arsenite for 30 min and subjected to polysome profiling. Ten fractions collected from the sucrose gradient were prepared for western blot analyses against pan-ADPr reagent. (E) First five fractions from cells expressing GFP or GFP-tagged nsP3 were run side by side and blot against pan-ADPr reagent to reveal ADP-ribosylation pattern.

To investigate the impact of nsP3 expression on ADP-ribosylation levels, we transfected cells with either GFP-vector or GFP-tagged nsP3, treated them with arsenite, and performed polysome profiling fractionation to enrich SG-associated proteins^9,10^ (Fig. 1D). The fractionated samples were then probed with a pan-ADP-ribose reagent to detect ADP-ribosylated proteins, irrespective of their forms. An accumulation of ADP-ribosylated proteins associated with fractions #2 to #6, which contain monosomes and untranslated mRNPs, in control cells expressing the GFP vector (Fig. 1D). In contrast, cells expressing GFP-tagged nsP3 exhibited a reduction in ADP-ribosylation signals in these fractions (Fig. 1D-E). Western blot analysis of SG-associated translation factor RACK1 was enriched in monosome-containing fractions, indicative of translation arrest, as expected from arsenite treatment^10^. These findings suggest that nsP3 expression leads to a decrease in ADP-ribosylation on target proteins, which may play a crucial role in inhibiting SG assembly.

### The MARylating enzyme PARP10 is critical for the initial stage of SG assembly

Given the strong inhibitory effect on SG assembly by MAR-degrading nsP3, we hypothesized that MARylation is critical for SG assembly. To pinpoint the specific mono-ADP-ribosyltransferase(s) involved, we systematically knocked down each enzyme and assessed their impact on SG formation after a 30-min arsenite treatment. As expected, cells transfected with the non-targeting siRNA control (siCon) exhibited SGs in ∼90% of cells, as indicated by SG components G3BP1 and eIF3b (Fig. 2A-B, S1A). The knockdown of most PARPs did not significantly affect SG assembly, except for PARP10 and PARP14. Their knockdown led to reduced SG assembly, with SGs present in 44% and 58% of cells, respectively (Fig 2A-B). Given the stronger inhibitory effect of PARP10, we subsequently focused on this mono-ADP-ribosyltransferase^26^.

**Figure 2.**
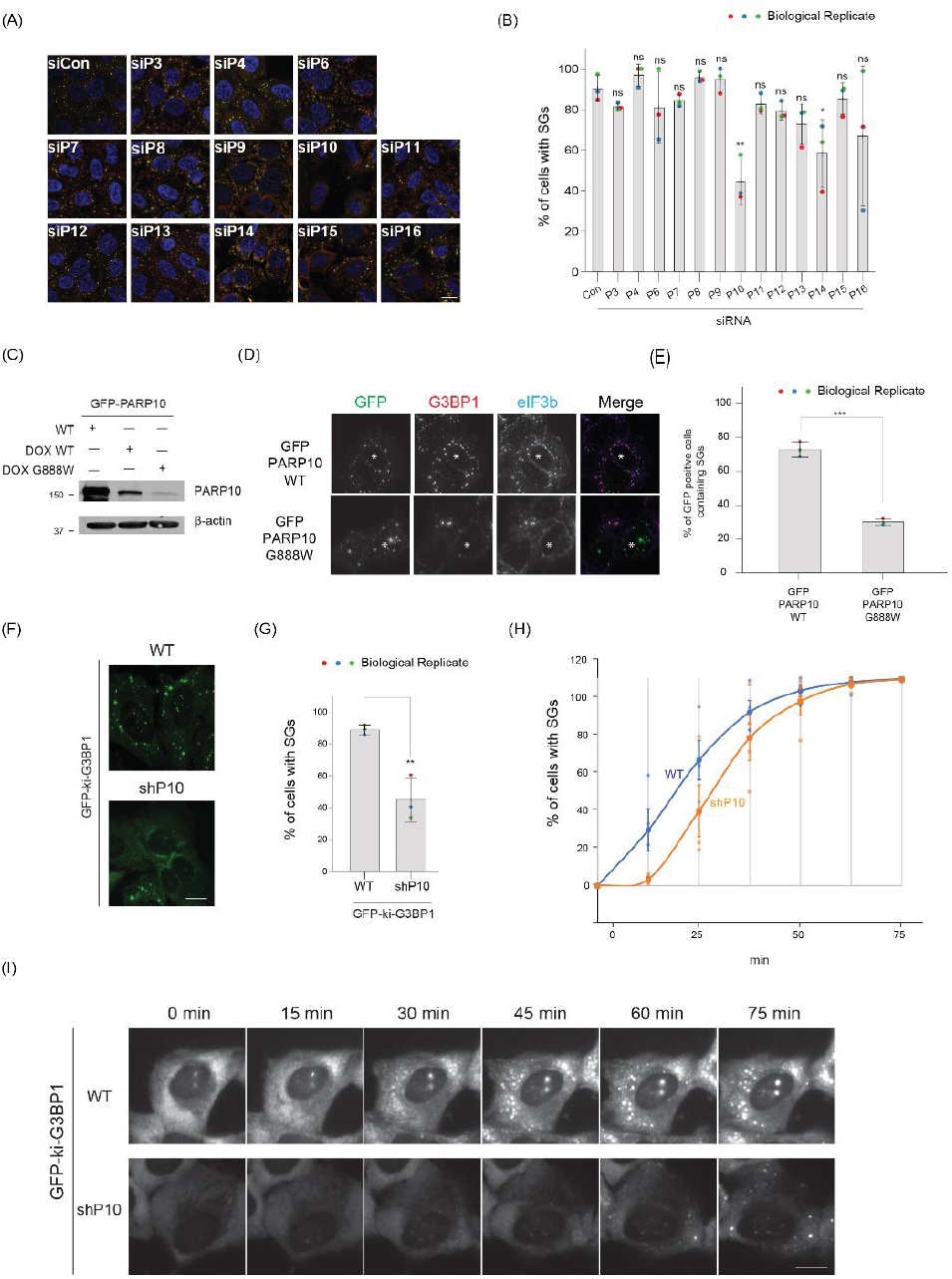
PARP10 is essential for SG assembly. (A) U2OS cells were transfected with siRNA control (siCon) or siRNA against different PARPs. 36 h after transfection, cells were treated with 0.2 mM arsenite for 30 min, fixed, and immunostained with G3BP1 (green) or eIF3b (red) antibodies. Scale bar, 10 μm. (B) Bar graph showing the percentage of cells with SGs in (A). (C) Expression level of constitutively active GFP-PARP10, dox-inducible GFP-tagged PARP10 WT and G888W mutant in U2OS cells. Dox was added for 24 h to induce the expression of PARP10 WT and G888W mutant. (D) Dox-inducible GFP-tagged PARP10 WT or G888W mutant constructs were transfected into U2OS cells. After 12 h, cells were changed to fresh medium containing Dox and incubated for 24 h at 37°C. Cells were then treated with 0.2 mM arsenite for 30 min and fixed for immunofluorescence. (E) Bar graph showing the percentage of cells containing SGs. (F) GFP-ki-G3BP1 WT or shPARP10 stable cell lines were treated with 0.2 mM arsenite for 30 min and fixed for immunofluorescence. (G) Bar graph showing the percentage of cells with SG in (F). (H) Graph showing the percentage of SG as a time series upon 0.2 mM arsenite treatment in WT and shPARP10 cell line. (I) Snapshot of live-cell imaging data from WT and shPARP10 cell line from (H). Asterisk indicates co-transfected cells. Scale bar, 10 μm.

The effect of PARP10 on SG assembly was confirmed with different siRNAs in U2OS cells and A549 cells (Fig. S1B-E). Taken together, our findings suggest that SG assembly requires PARP10 and, to a lesser degree, PARP14.

Previously, we identified the localization of PARPs 5a, 12, 13, 14, and 15 at SGs, but not PARP10^14^. Given the pronounced effect of PARP10 knockdown on SG assembly, we re-examined its relationship with SGs. As reliable antibodies for staining endogenous PARP10 were lacking, we utilized GFP-tagged PARP10 to observe its cellular localization. Cells expressing PARP10 formed condensates distinct from SGs (Fig. S2A)^27^. Additionally, higher PARP10 levels in cells inhibited SG formation, consistent with previous findings that overexpressing PARP10 depletes NAD^+^, potentially hindering ADP-ribosylation critical for SG assembly^28^.

To address this limitation, we employed a doxycycline-based system to induce a lower PARP10 expression (Fig. 2C-E). This reduced expression did not affect SG assembly, with 87% of cells exhibiting SGs upon arsenite treatment, similar to untransfected cells. However, even at these expression levels, PARP10 remained in condensates distinct from SGs. We then explored whether this unique localization is tied to catalytic activity. Using the PARP10 catalytic dead mutant G888W^26^, we observed that this mutant was still localized as condensates (Fig. 2D). Intriguingly, its presence in cells significantly inhibited SG assembly, with only 36% of cells with SGs (Fig. 2E), suggesting that PARP10 enzymatic activity impacts SG assembly.

As is common in the field, we measured the impact of gene knockdown or overexpression on SG assembly at a fixed time point^10,23^. However, endpoint-based assays only provide a snapshot and might not capture the entire dynamics of SG assembly. To investigate whether PARP10 impacts the kinetics of SG assembly, we generated a cell line expressing shRNA targeting PARP10 (shPARP10) and performed live cell imaging. Specifically, lentiviral transduction introduced shRNAs into a U2OS cell line that is endogenously tagged with the SG core component G3BP1 with GFP (GFP-ki-G3BP1; Fig. 2F). Consistent with the transient knockdown results, the shPARP10 line showed significant inhibition of SG assembly (45% cells with SGs) compared to the parental line (89%) 30 min post arsenite treatment (Fig. 2F, G, S2B). Live-cell imaging further revealed that SG assembly was notably delayed in the shPARP10 cell line relative to the parental line (Fig. 2H, I). However, as time progressed, SG assembly in both lines reached the same plateau at ∼75 min (Fig. 2H). These findings collectively underscore the crucial role of PARP10 in the initial stages of SG assembly.

### PARP10 is required for G3BP1-mediated SG assembly

G3BP1 and its paralog G3BP2 are essential SG components^23,29^. The G3BP1/2 double knockout (dKO) cell line cannot form SGs upon various stress conditions, including arsenite treatment. However, reintroducing at least one of the G3BPs (either G3BP1 or G3BP2) restores SG formation in the dKO cell line^29^. To delve deeper into the role of PARP10 in G3BP1-mediated SG assembly, we transfected the G3BP1/2 dKO cell line with either PARP10 siRNA or non-targeting siRNA (siCon), then subsequently transfected it with GFP-G3BP1 (Fig. 3A, B). The overexpression of GFP-G3BP1 restored SG formation in ∼55% of cells, as evidenced by the presence of eIF3b in GFP-G3BP1 condensates (Fig. 3C, D). In contrast, only ∼9% of cells with PARP10 knockdown exhibited SGs. These data suggest PARP10 is required for G3BP1-mediated SG assembly.

**Figure 3.**
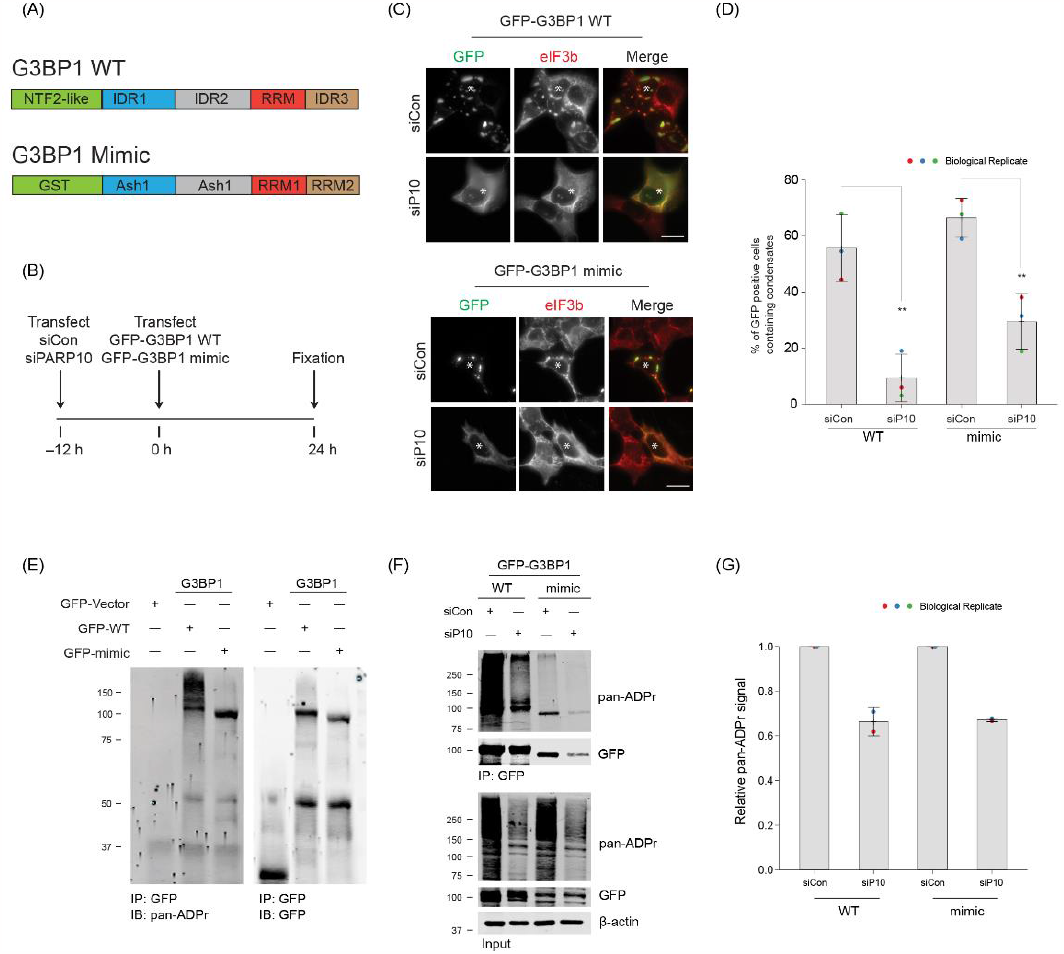
PARP10-mediated ADP-ribosylation is critical for SG condensation. (A) Domain structure of G3BP1 WT and mimic construct. (B) Experimental flow for panels C and F. (C) G3BP1/2 dKO cells were either treated with siCon or siPARP10 for 24 h. After 24 h, cells were transfected with either GFP-tagged G3BP1 WT (Top panel) or synthetic G3BP1 construct (Bottom panel) for an additional 24 h. Post transfection, cells were fixed and immunostained against anti-eIF3b antibodies. Asterisk indicates co-transfected cells. Scale bar, 10 μm. (D) Bar graph showing the percentage of GFP-positive cells containing SGs. (E) G3BP1/2 dKO cells were transfected with either GFP, GFP-tagged G3BP1 WT or synthetic G3BP1 construct for 24 h. Post transfection, cells were immunoprecipitated using anti-GFP antibodies, and blotted against pan-ADP ribose reagent. (F) G3BP1/2 dKO cells were either treated with siCon or siPARP10 for 24 h, followed by transfection with either GFP-tagged G3BP1 WT or synthetic construct for 48 h. Post transfection, cells were lysed, immunoprecipitated, and blotted against pan-ADP ribose reagent. (G) Quantification of pan-ADP-ribose band intensity between siCon and siPARP10 conditions.

Building on this data, we explored a simplistic model—G3BP1 ADP-ribosylation, facilitated by PARP10, drives SG assembly. Supporting this model, our prior studies show that G3BP1 ADP-ribosylation is strongly correlated with SG assembly: G3BP1 ADP-ribosylation increases under conditions when stress granules form, whereas this ADP-ribosylation reduces upon infection when the viral-encoded, MAR-degrading nsP3 macrodomain inhibits SG assembly^14,18^. Indeed, when G3BP1 was reintroduced into the G3BP1/2 dKO cell line, leading to SG restoration, G3BP1 was found to be ADP-ribosylated (Fig. 3E).

However, this ADP-ribosylation was significantly reduced upon PARP10 knockdown (Fig. 3F, G). This data underscores the pivotal role of PARP10 in the ADP-ribosylation of G3BP1.

A recent study indicates that SG formation in G3BP1/2 dKO cells can be restored not only by reintroducing G3BP1 but also by using a construct mimicking G3BP1^29^. This mimic replaces G3BP1’s essential domains responsible for SG assembly with analogous domains from other proteins (Fig. 3A). To investigate if PARP10 regulates SG assembly even without G3BP1, we examined the effect of PARP10 knockdown on SG formation using this mimic. Much like reintroducing wild-type G3BP1, expressing the GFP-tagged mimic led to SG restoration in ∼66% of the G3BP1/2 dKO cells, as verified by the colocalization of eIF3b and GFP signals (Fig. 3C, D). Yet, with PARP10 knockdown, the percentage of cells exhibiting SGs with the G3BP1 mimic dropped to 29% (Fig. 3C, D). As in the case of wild-type G3BP1, its mimic was also ADP-ribosylated in a PARP10-dependent manner (Fig. 3E, F). Taken together, these data suggest that PARP10 regulates SG assembly, regardless of whether the assembly is driven by G3BP1 or its modeled mechanism.

### SG core composition is altered upon PARP10 knockdown

While previous studies have underscored the significance of PAR in forming SGs by recruiting proteins through non-covalent interactions^14,20–22^, the involvement of PARP10—which adds MAR and not PAR^26^—presents a conundrum. To probe deeper into this relationship, we analyzed the ADP-ribosylation form of G3BP1. By using antibodies that specifically detect either MAR or PAR^30,31^, we probed the GFP immunoprecipitates from dKO cells expressing GFP-G3BP1. As controls, PARP1 and PARP10, known to be PARylated and MARylated, were included respectively^17^. Unexpectedly, our data indicated that G3BP1 is PARylated, not MARylated (Fig. 4A). Since ADP-ribosylation of G3BP1 is dependent on PARP10, it is plausible that PARP10 initiates the ADP-ribosylation of G3BP1, which can then be further PARylated by other enzymes.

**Figure 4.**
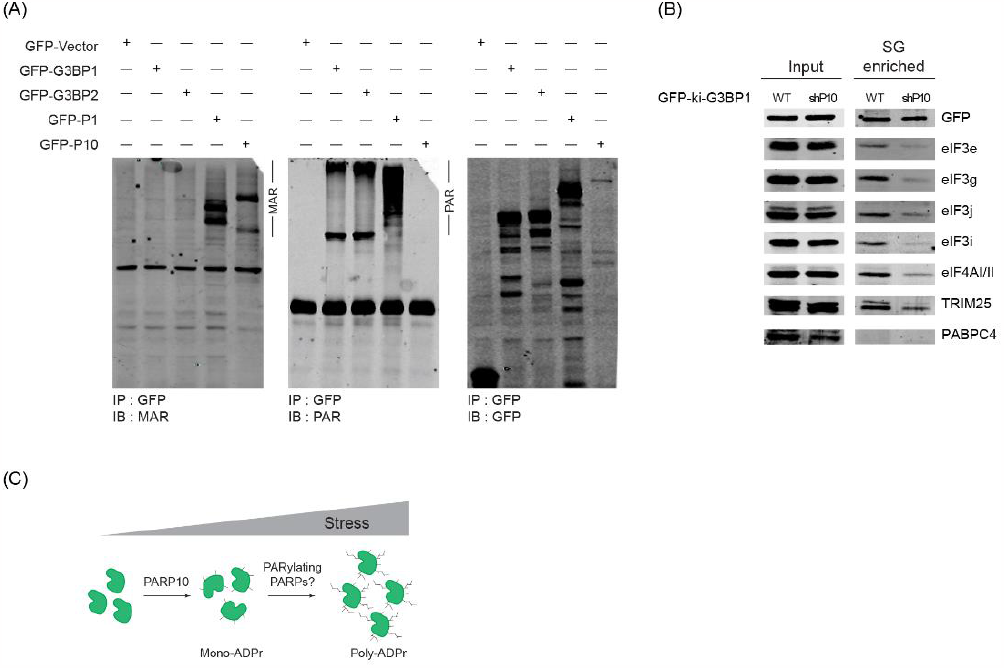
PARP10 initiates SG condensation and recruits SG components. (A) G3BP1/2 dKO cells were transfected with vectors encoding GFP, GFP-tagged G3BP1, G3BP2, PARP1, or PARP10 after 24 hours, immunoprecipitated using GFP-antibodies, and subjected to western blotting against antibodies specific to MAR or PAR. (B) GFP-ki-G3BP1 WT and shPARP10 cells were treated with 0.2 mM arsenite for 30 min. Following stress induction, SGs were isolated, and subjected to western blot analyses against indicated antibodies. (C) Proposed model of PARP10-mediated G3BP1 MARylation and further extension of ADP-ribose chain by other PARPs.

Considering the critical role of PAR in recruiting proteins to SGs, we hypothesized that reduced ADP-ribosylation due to PARP10 knockdown might alter SG composition. To test this hypothesis, we employed a differential centrifugation method to isolate SG cores^32^. We then probed these cores from different conditions with antibodies against several SG components^33^. Given that several translation factors, such as eIF3g, bind to PAR^18^, we particularly focused on this class of SG components. Intriguingly, SG cores from the shPARP10 cell line exhibited an altered composition. Translation factors eIF3g, eIF3j, and eIF4AI/II were less enriched compared to those from the parental line (Fig. 4B). Collectively, our data suggest that PARP10 plays a pivotal role in SG assembly by initiating ADP-ribosylation of G3BP1, which then evolves to form PAR, thereby recruiting additional SG components.

## DISCUSSION

Here, we pinpointed MARylation as a critical form of ADP-ribosylation integral to SG assembly. The enzyme PARP10, responsible for MARylation, regulates the initial stage of SG assembly and targets the core component G3BP1. A decrease in PARP10 reduced the levels of translation factors within the SG core. Additionally, unexpected observations emerged from our study, which are discussed below.

### MARylating Enzymes Drives PAR-enriched Stress Granule Assembly

Many SG proteins are PARylated or bind PAR^14,19–21^. Overexpressing PARG suppresses SG assembly, while its knockdown impedes disassembly, indicating the critical role of PAR in maintaining SG integrity. However, inhibitors against PAR-adding PARPs, including the SG-localized PARP5a, do not suppress SG formation. Instead, these inhibitors against PARP1/2 or PARP5a/b only block the localization of select components to SG, suggesting that PAR controls SG composition^22^. On the other hand, MARylation emerges as a key player in SG assembly. Three out of five PARPs localized in SGs are MAR-adding, and overexpression of any of them is sufficient to induce SG formation. Our systematic PARP knockdown now further revealed MAR-adding PARP10 and, to a lesser extent, PARP14 as critical for SG formation. Unlike PARP14, PARP10 is not enriched in SGs, suggesting that the MARylation responsible for SG formation can be initiated outside SGs.

Intriguingly, PARP10 levels, and presumably associated ADP-ribosylation activity, must be tightly regulated to ensure proper SG formation. Its overexpression suppressed SG formation, whereas low expression did not. Yet, even at low levels, a catalytically dead PARP10 mutant inhibited SG formation in the presence of endogenous PARP10. These data suggest a dominant-negative effect, where the mutant interferes with the function of the wild-type PARP10 or other essential components of SG formation. One potential mechanism is that either an excess of PARP10 or its lack of catalytic activity might sequester key substrates or cofactors required for SG formation.

### The Role of PARP10 and Its Substrates in Stress Granule Assembly

G3BP1 is central to SG assembly, undergoing significant ADP-ribosylation upon stress. In this study, we identified G3BP1 as a substrate for PARP10. In G3BP1/2 dKO cells, reintroducing G3BP1 led to its ADP-ribosylation and restoration of SGs—both processes dependent on PARP10. Therefore, while G3BP1 is essential for SG assembly, its function may depend on PARP10. Interestingly, a construct that mimics the domain architecture of G3BP1 also required PARP10 to restore SGs in G3BP1 dKO cells^29,34^. This finding raises the intriguing possibility that ADP-ribosylation of any core SG component could be sufficient to initiate SG assembly, irrespective of its identity. Alternatively, the mimic may emulate the native G3BP1’s function, recruiting specific proteins for PARP10 modification. Future studies should determine if SG assembly is primarily driven by ADP-ribosylation of core SG components or on specific proteins that the core components recruit. Given that ADP-ribosylation is observed in multiple SG proteins, including TIA-1, TDP-43, and hnRNPA1^21,22^, additional studies should examine their potential involvement in SG assembly.

### Working Model: MARylation as a rate-limiting step of SG assembly

ADP-ribosylation is instrumental in the initial SG assembly kinetics. Previous findings show that the FDA-approved PARP1/2 inhibitor, Olaparib, diminishes the number of cells with SGs during the early stages, with no discernible differences in later stages^22^. This trend is echoed in our live-cell imaging of the shPARP10 cell line, which displays a delay in SG assembly compared to the parental line. Moreover, the marked inhibitory effect of the viral MAR-degrading nsP3, especially when contrasted with the human PAR-degrading PARG, emphasizes the pivotal role of MARylation in regulating SG assembly dynamics^18^.

Building on these observations, we propose that MARylation of one or more substrates is crucial for SG assembly. Data from PARP10 or PARP14 knockdowns points towards potential redundancy in the MAR-adding mechanism. These MARylated substrates likely act as anchors, facilitating protein interactions that initiate SG assembly. Moreover, PAR-adding PARP5a or PARP1 might extend this MARylation to PARylated forms for additional protein interactions through this polynucleotide-like polymer. In line with this model, we identified PARP10-dependent modification of G3BP1 as PARylation, rather than MARylation.

One scenario is that PARP10 mediates the initial stage of assembly, where ADP-ribosylation of G3BP1 and its inherent RNA and protein-protein interactions drive the condensation process (Fig. 4C)^23,35^.Consistent with prior studies, G3BP1 predominantly interacts with the same protein partners even in the unstressed state^36^. However, upon SG formation, the concentration of G3BP1 and these interacting proteins is likely to increase^33,36^. We postulated that ADP-ribosylation, in addition to RNA, enhances G3BP1’s interactions with proteins via PAR. This positions ADP-ribosylated G3BP1 as a super-scaffold within SGs^18,19,21,22^, bridging proteins through interactions with two nucleic acid polymers. The reduced presence of some PAR-binding translation factors in SG cores following PARP10 knockdown further supports this possibility. A comprehensive SG proteome comparison between wild-type and PARP10 knockdown cells may yield deeper insights.

From both the perspective of ADP-ribosylation addition and its degradation, we identified pivotal roles for MAR-adding and MAR-degrading enzymes in the assembly of RNA-enriched SGs. Therefore, we propose that MARylation acts as the rate-limiting step for SG assembly, possibly by initiating the PAR chain. Supporting this idea, our recent work demonstrates that even a 1:1000 sub-stoichiometric ratio of PAR can potently condense the SG-localized RNA-binding protein FUS^37^. While FUS co-binds and co-condenses with RNA and PAR, RNA can also enter pre-formed FUS-PAR condensates, displacing a majority of the existing PAR^38^. Notably, both PARP10 and PARP14 possess RNA-binding domains, hinting at a deeper interplay with RNA dynamics in SG assembly. These findings, along with the kinetic role established for ADP-ribosylation in SG assembly, suggest a model: Initiated by rate-limiting MARylation, PAR triggers SG protein condensation. These early-stage transient PAR-protein interactions eventually transition to more stable interactions between proteins and RNA, leading to SG formation.

### Limitations

Our data point to PARP10 as the primary PARP involved in SG assembly. However, it is possible that different cell lines might depend more on other MAR-adding enzymes like PARP14^39^. While the effects of the catalytic dead mutant highlight the significance of PARP10 activity, we cannot conclusively determine if the observed knockdown effects are due to its physical absence or its enzymatic function. Experiments with a PARP10 inhibitor could provide more clarity. While our study is centered on SG assembly, the significance of MARylation or PARylation for SG disassembly remains unclear. Our previous data indicate that knocking down PARG, resulting in increased PAR levels, prolongs SG disassembly. Conversely, the catalytic mutant of the viral MAR-degrading enzyme delays SG disassembly during infection. Exploring disassembly further could reveal novel therapeutic strategies targeting SGs or SG-like condensates in viral infection, cancer, and neurodegeneration.

## MATERIALS AND METHODS

### Cell Culture, Chemicals, and Transfection

U2OS and A549 cells were obtained from the American Type Culture Collection (ATCC). Cells were maintained in Dulbecco’s modified Eagle medium (DMEM) (Gibco) containing 10% heat-inactivated fetal bovine serum (FBS) (Gibco, Life Technologies). Plasmids were transfected using JetPrime from Polyplus as per the manufacturer’s protocol.

### Immunoblot Analysis

Cells were lysed in RIPA buffer (50 mM Tris-Cl pH 8.0, 150 mM NaCl, 0.1% sodium dodecyl sulfate [SDS], 1% Nonidet P-40, 1 mM EDTA with phosphatase and protease inhibitor cocktails) with 10 μM Olaparib (PARP inhibitor) and 10 μM PDD 00017273 (PARG inhibitor) on ice for 15 min, followed by centrifugation at 14,000 rpm for 15 min at 4°C. Protein samples were acetone-precipitated for at least 1 h at −20°C. Precipitates were centrifuged at 13,000 rpm, 4 °C for 15 min, and the air-dried pellets were then diluted in 1×SDS sample buffer. The samples were resolved in polyacrylamide gel electrophoresis and blotted with appropriate primary antibodies.

### Immunofluorescence

U2OS cells (∼6×10^4^) grown on coverslips were treated with the 0.2 mM arsenite for 30 min. Post-treatment, cells were washed twice with 1×PBS, fixed with 4% paraformaldehyde for 15 min at room temperature (RT), permeabilized with ice-cold methanol for 10 min, and washed twice with 1×PBS. Cells were then blocked with 5% normal horse serum in 1×PBS containing 0.02% sodium azide for 1 h at RT. All primary antibodies were diluted in blocking buffer and incubated either 1 h at RT or overnight at 4 °C, followed by three washes with 1×PBS, 10 min each. Finally, cells were incubated with appropriate secondary antibodies diluted in blocking buffer (1:500) for 1 h at RT, washed three times with 1×PBS, and the coverslips were mounted on glass slides using Prolong Gold.

### Immunoprecipitation

U2OS wild-type or G3BP1/2 dKO cells were lysed in cold lysis buffer (CLB) (50 mM HEPES pH 7.4, 150 mM NaCl, 1 mM MgCl_2_, 1 mM ethylene glycol-bis(β-aminoethyl ether)-N,N,N′,N′-tetraacetic acid, 1% Triton X-100 supplemented with 1 mM NaF, 1 mM PMSF, 10 μM Olaparib and 10 μM PDD 00017273. Cell lysates were mixed at 4°C for 15 min, spun down for 15 min at 13,000 rpm, and the supernatant fluid was collected in a new tube. GFP-magnetic bead complex was prepared by incubating anti-GFP (3E6, Invitrogen) with DYNA magnetic beads (10004D, Invitrogen) at room temperature for 10 min. The complex was then washed once with lysis buffer and the cleared lysates were added. Samples were incubated for 2 h at 4°C. Beads were washed once with CLB, twice with high-salt CLB (300 mM NaCl), followed by a final wash with CLB. The precipitates were then boiled with 1×SDS sample buffer for 10 min at∼85 °C. All experiments were performed at least thrice.

### Live-cell imaging

GFP-ki-G3BP1 WT and shPARP10 cells were seeded in 2-well chambered cover glass (Nunc Lab-Tek) at around 70% confluency. One hour before imaging, cells were pre-incubated with CO_2-_ independent medium containing 20% FBS. Cells were incubated with 0.2 mM arsenite and live-cell imaging was performed simultaneously for WT and shPARP10 cell lines. A total of 15–20 random fields were chosen and imaged, and the presence of SG was monitored as microscopically visible condensates in the FITC channel. Images were acquired at 5-min intervals for 90 min, controlled by the SoftWorx suite (GE Healthcare). Cells were imaged using a DeltaVision Elite system (GE Healthcare) microscope equipped with ×40 (1.516 N.A. oil) immersion objectives, a high-speed CCD Camera (Cool SNAP HQ2), appropriate filter sets for FITC, and an incubation chamber (37 °C and 80% humidity).

### Isolation of stress granules

Stress granule isolation was performed according to a previously described method^32,33^. Briefly, GFP-ki-G3BP1 or GFP-ki-shPARP10 cell lines in 15 cm tissue-culture dishes (6 plates per condition) were treated with 0.2 mM sodium arsenite for 30 min. After treatment, cells were washed once with cold PBS, scraped in PBS, and flash frozen with liquid N_2_ and the cell pellets were stored at −80°C. The pellets were thawed on ice for 5 min and re-suspended in 200 μl stress granule lysis buffer (50 mM Tris-HCl, pH 7.4, 100 mM potassium acetate, 2 mM magnesium acetate, 0.5 mM DTT, 50 μg/mL heparin, 0.5% NP40, 1:5000 Antifoam B, 1x EDTA-free protease inhibitor cocktail and 1:500 RNase inhibitor). To efficiently lyse the cells, 10 passages through the 25G gauge 5/8 syringe were performed. After lysis, spin the samples at 1000 x g for 5 min at 4°C to pellet cell debris, and the supernatants were transferred to new tubes and centrifuged at 18,000 x g for 20 min at 4°C. The supernatants were discarded, and the pellets were resuspended with 1 ml stress granule lysis buffer.

The samples were again centrifuged at 18,000 x g for 20 min at 4°C. The pellets were re-suspended in 60 μl stress granule lysis buffer followed by spin at 850 x g for 2 min at 4°C, and the supernatants representing the stress granule enriched fractions were boiled with 1x SDS sample buffer for 10 min at ∼90°C.

### Stress granule quantification

We classified cells as SG-negative if the entire cell lacked both G3BP1 and eIF3b condensates. For quantitation, random 40× fields were chosen and a total of between 80 and 120 cells were counted per condition. For quantification of SGs from the live-cell imaging experiments, the number of SGs present at each time was calculated and plotted. In experiments involving the G3BP1/2 dKO cell line, cells were scored as SG positive when the GFP-expressing cells contained eIF3b condensate as well. All experiments were repeated three independent times.

### Statistical Analysis

Data were presented as mean ± SD and groups were compared using two-tailed unpaired Student’s t-test and Kolmogorov–Smirnov test. P< 0.05 was considered statistically significant.

## Author Contribution

A.K.J. and A.K.L.L. designed the research; A.K.J. and K.B. performed the research; and A.K.J. and A.K.L.L. wrote the paper.

## Acknowledgement

We thank members of the A.K.L.L. laboratory for their critiques of the manuscript. We thank Dr. Nancy Kedersha and Dr. Paul Anderson for G3BP1/2 dKO cells and G3BP1 constructs, and Dr. Paul Taylor for GFP-ki-G3BP1 cell line. The authors would also like to thank Dr. Lee Kraus and colleagues for sharing their related paper prior to their submission. This work was supported by a Johns Hopkins Sol Goldman pancreatic cancer research center pilot project grant (A.K.L.L.) and NIH grant R01GM104135 (A.K.L.L.)

## Resource table

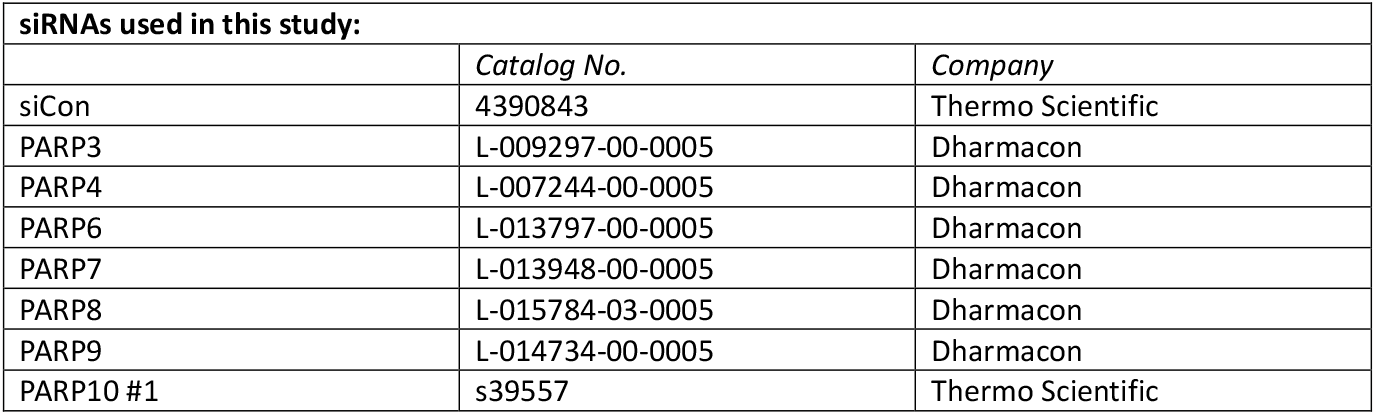
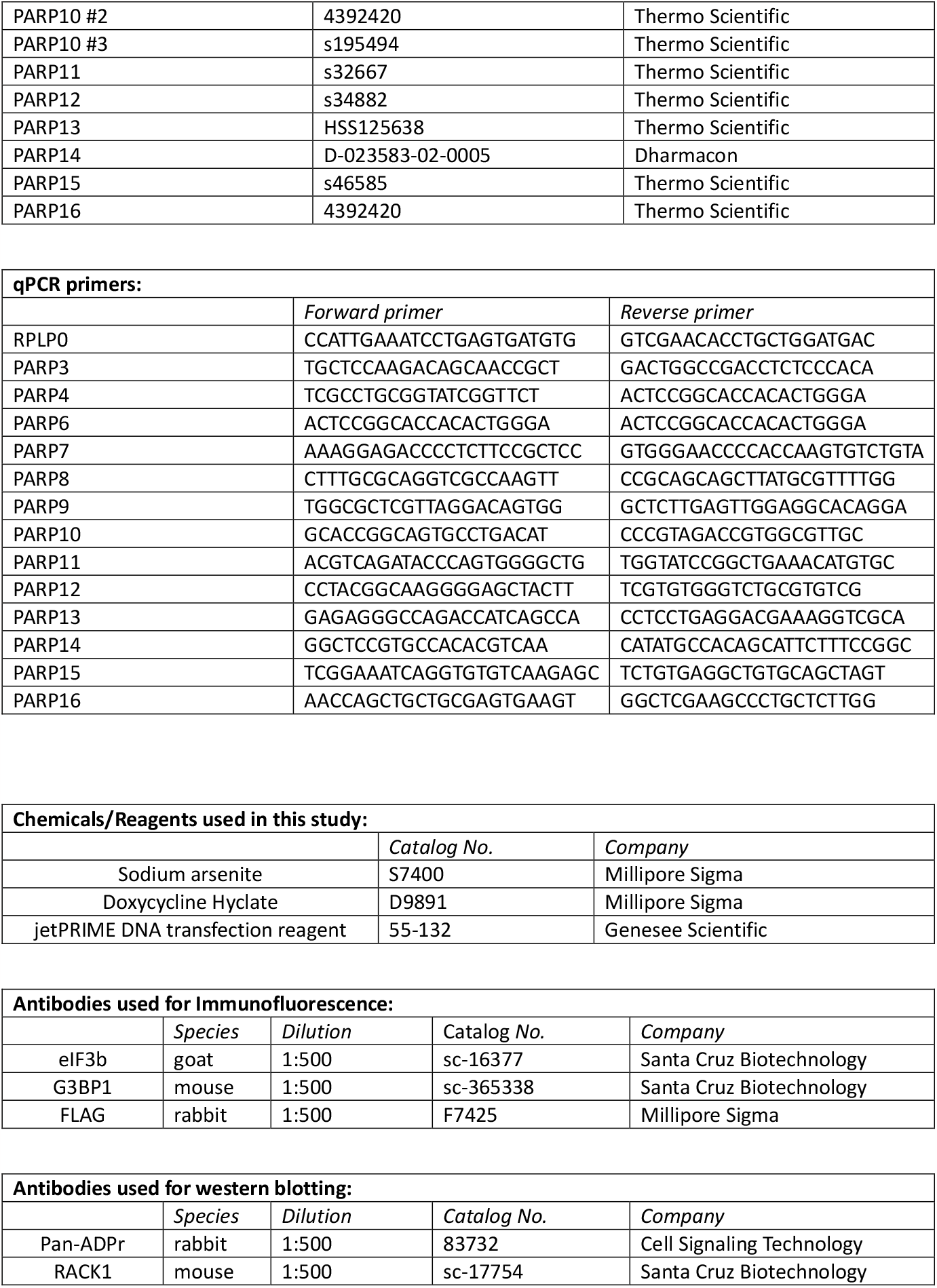
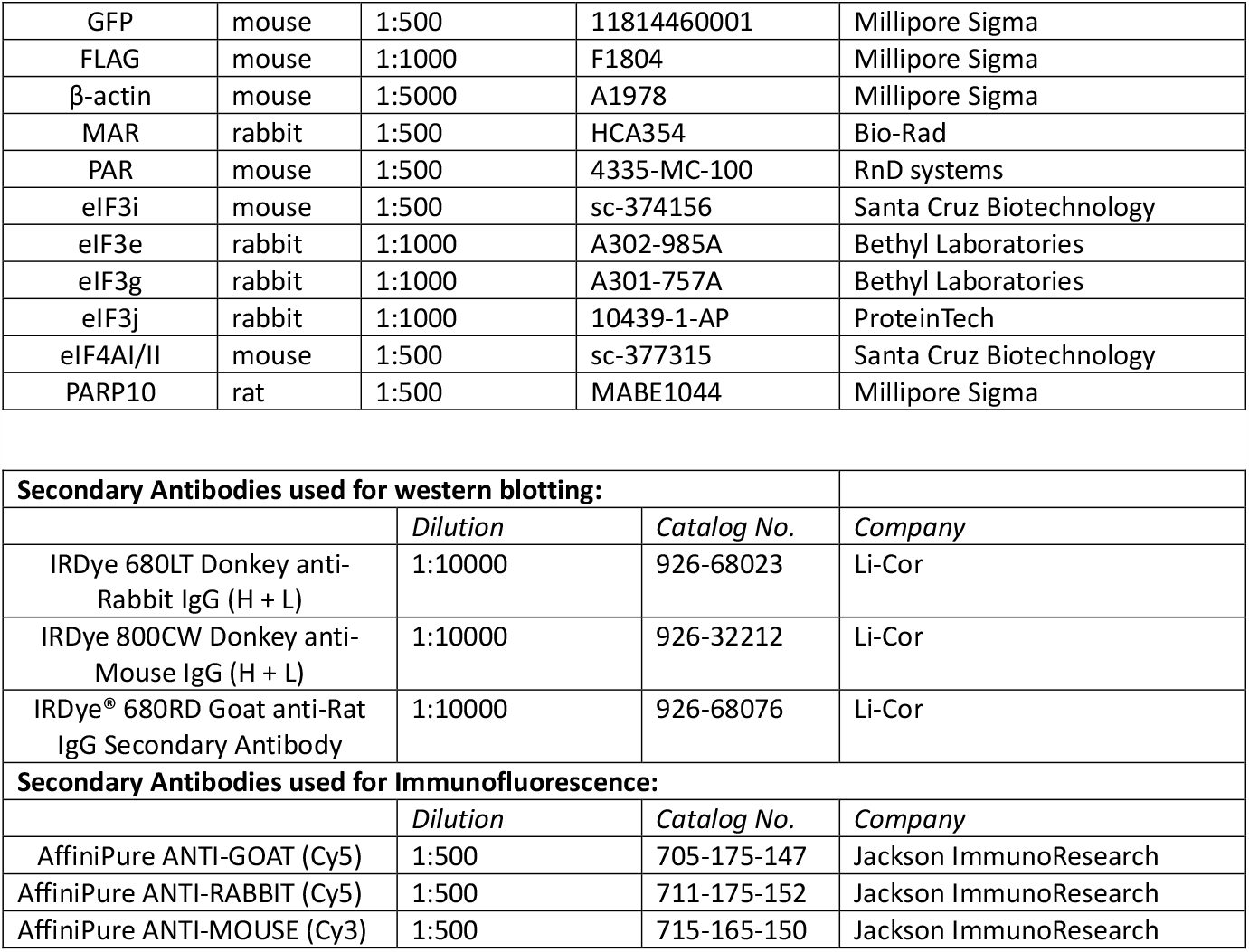

