## Supplemental Figures for "PARP10 is Critical for Stress Granule Initiation"

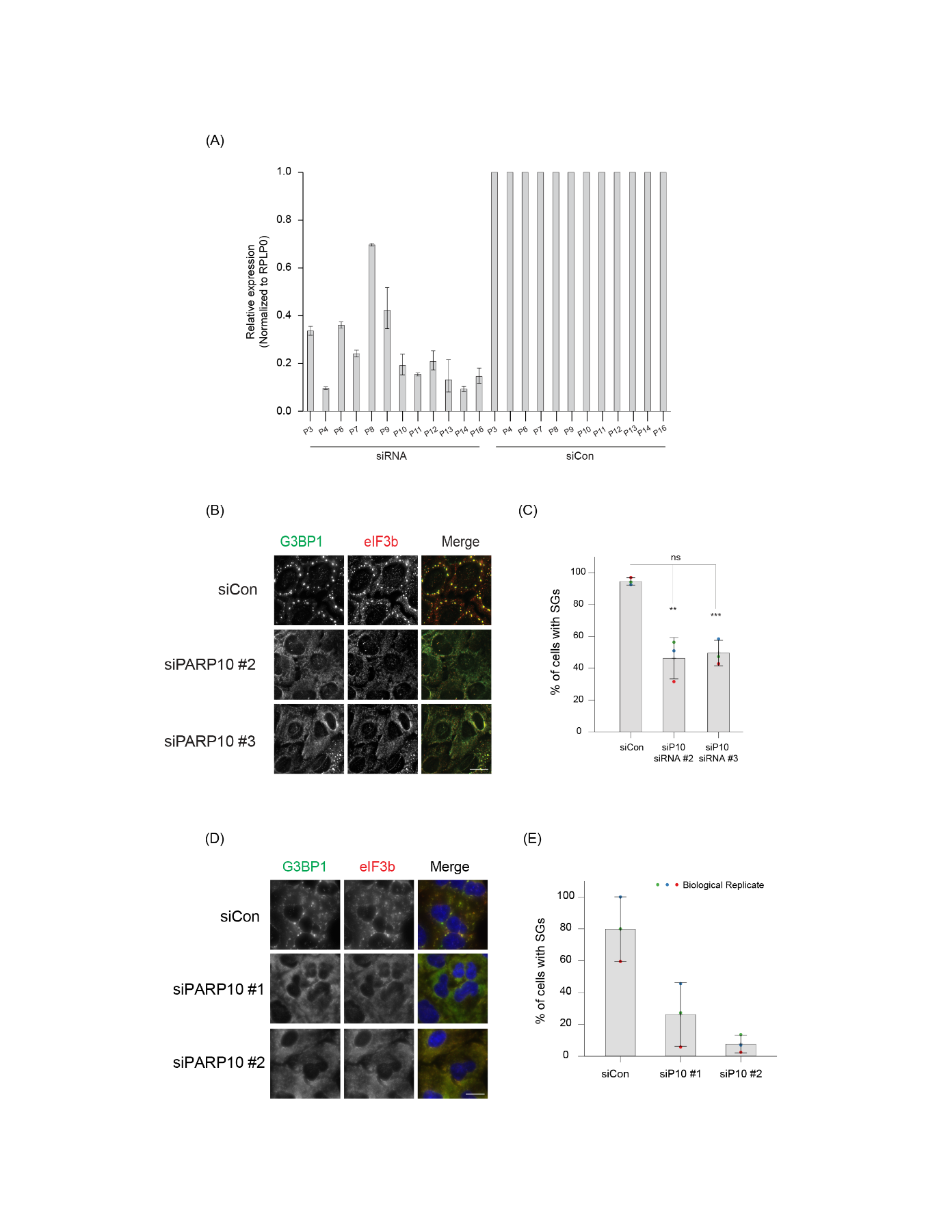
**Supplementary Figures:**

**Supplementary Figure 1.** (A) Bar graph showing the knockdown efficiency of different PARPs in Fig. 2A. (B) U2OS cells were transfected with siCon or siRNAs targeting different regions of PARP10 (#2 and #3) for 36 h. After 36 h, cells were treated with 0.2 mM arsenite for 30 min, fixed, and immunostained against anti-G3BP1 and anti-eIF3b antibodies. (C) Bar graph showing the percentage of cells with SGs in (B). (D) Fig. S2B was repeated in A549 cells. (E) Bar graph showing the percentage of cells containing SGs as in Fig. S1D.

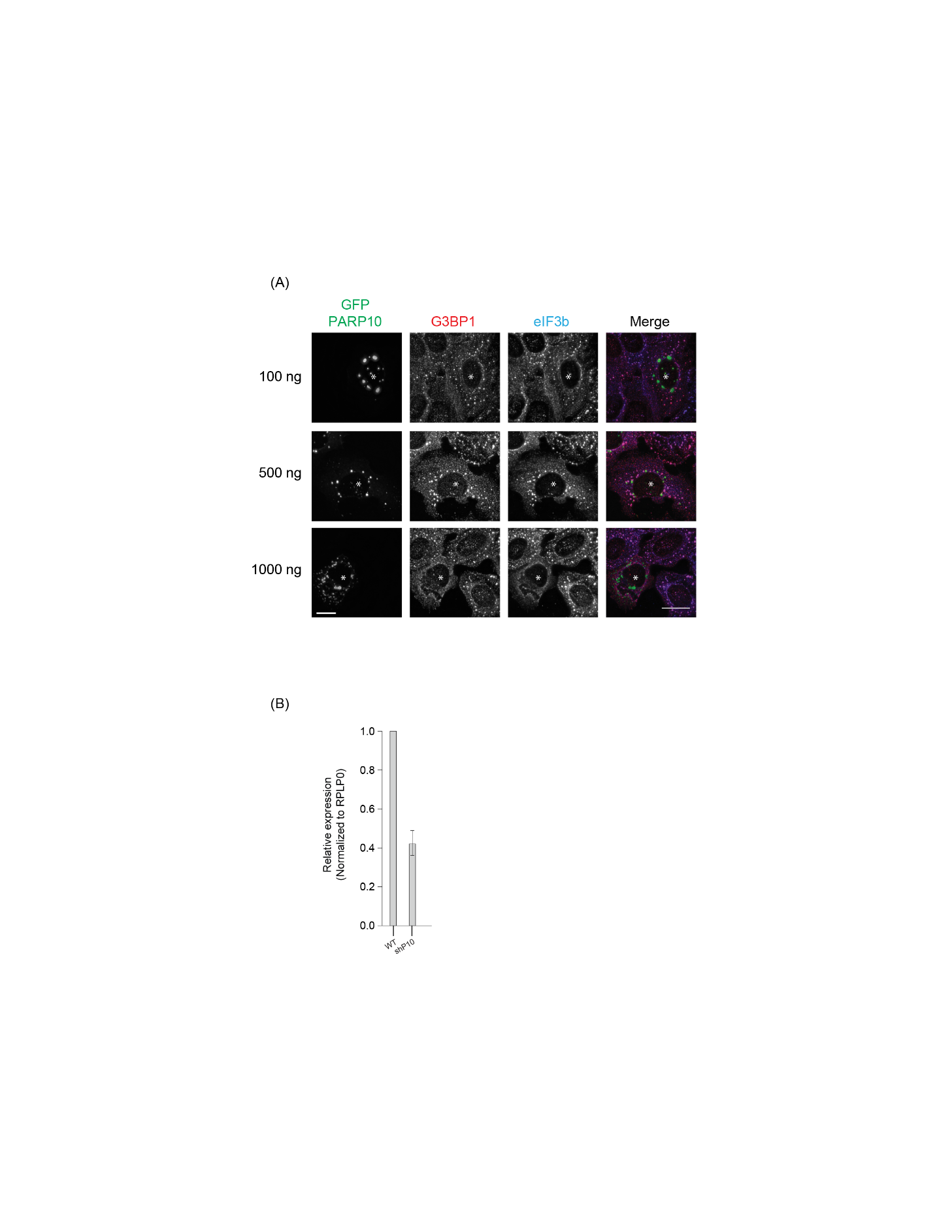

**Supplementary Figure 2.** (A) GFP-tagged PARP10 construct was transfected into U2OS cells at different concentrations (100 ng, 500 ng, 1000 ng). 24 hour post transfection, cells were treated with 0.2 mM arsenite for 30 min, fixed and immunostained against anti-eIF3b and G3BP1 antibodies. (B) Knockdown efficiency of PARP10 in shPARP10 cell line as in Fig. 2C.
